# Role of the circadian clock “Death-Loop” in the DNA damage response underpinning cancer treatment resistance

**DOI:** 10.1101/2022.01.14.476363

**Authors:** Ninel Miriam Vainshelbaum, Kristine Salmina, Bogdan I. Gerashchenko, Marija Lazovska, Pawel Zayakin, Mark S. Cragg, Dace Pjanova, Jekaterina Erenpreisa

## Abstract

The Circadian Clock (CC) drives the normal cell cycle and reciprocally regulates telomere elongation. However, it can be deregulated in cancer, embryonic stem cells (ESC), and the early embryo. Here, its role in the resistance of cancer cells to genotoxic treatments was assessed in relation to whole-genome duplication (WGD) and telomere regulation. We first evaluated the DNA damage response of polyploid cancer cells and observed a similar impact on the cell cycle to that seen in ESC - overcoming G1/S, adapting DNA damage checkpoints, tolerating DNA damage, and coupling telomere erosion to accelerated cell senescence, favouring transition by mitotic slippage into the ploidy cycle (reversible polyploidy). Next, we revealed a positive correlation between cancer WGD and deregulation of CC assessed by bioinformatics on 11 primary cancer datasets (rho=0.83; p<0.01). As previously shown, the cancer cells undergoing mitotic slippage cast off telomere fragments with TERT, restore the telomeres by recombination and return their depolyploidised mitotic offspring to TERT-dependent telomere regulation. Through depolyploidisation and the CC “death loop” the telomeres and Hayflick limit count are thus again renewed. This mechanism along with similar inactivity of the CC in early embryos support a life-cycle (embryonic) concept of cancer.

## 1. Introduction: Whole-genome duplications (WGDs) induced in cancer radio- and chemoresistance reciprocally connect the mitotic and ploidy cycles coupling senescence to stemness

Malignant tumours are characterized by various degrees of aneu-polyploidy emerging from whole-genome duplications (WGD) that appear early in cancer evolution, progress with disease aggression, and correlate with resistance to anti-cancer treatments [1–6]. The first-line anticancer therapies (ionizing radiation and genotoxic drugs) kill most tumour cells in the first days after administration. However, these cells can also evoke transient polyploidy that can give rise to clonogenic para-diploid survivors several weeks or months after treatment cessation, recovering mitotic cycling, which disseminates, cause metastases, and repeatedly polyploidize with disease relapse [7–14]. Both in intrinsic and therapy-induced polyploidy, it appears that the two reproductive cycles, the rapid mitotic cycle and slow reversible polyploidy cycle, drive cancer cell immortality - their inter-relationship termed by us the “cancer cell life-cycle” [15–17]. The induction or enhancement of reversible polyploidy does not only assign the advantage of the genome multiplication masking lethal mutations and providing an option for more effective DNA repair [4,18–20] - in cancer, it is accompanied by a crucial change of cell biology - the reprogramming of somatic cells to a state of embryonal stemness [9,14,21–24]. This transition to stemness-related polyploidy is paradoxically coupled to, and even dependent on, accelerated cellular senescence (ACS) - first described as irreversible growth arrest in response to oncogenes, genotoxic and oxidative stress [25] signalling persistent DNA damage of compromised telomeres [26,27]. In cancer, both opposing phenomena, i.e. senescence and stemness, are intracellularly and paracrinally interacting, including immune cross-talk [28,29]; akin to the process of wound healing [30]. However, the relationship between this paradoxical pairing with reversible polyploidy [1,31–36] in the formation and metastasis of tumours [37] is poorly understood and often overlooked. Its mechanism appears to be based on an oscillation between the opposites: between senescence and stemness regulators in one plane, and between mitotic and polyploidy cycles on another [1,15]. To better understand the mechanics of this process we evaluated here the regulation of the cell cycle and DNA damage checkpoints in embryonic (ESC) and cancer stem cells (CSC). We also evaluated the potential role of the main driver of oscillatory cellular processes, the circadian clock (CC) in the DNA damage response and telomere maintenance, finally assessing the association of CC deregulation and WGD in primary cancers.

## 2. Regulation of the normal cell cycle and DNA damage checkpoints

The normal mitotic cell cycle consists of the G1, S, G2 and M phases. Their route and change is driven by corresponding cyclin-kinases. If DNA damage has occurred, cells can activate the G_1_, intra-S, and G_2_/M checkpoints and arrest the cell cycle to repair the damage. There are two major DNA damage signalling pathways - regulated by ATM/CHK2 and ATR/CHK1. The ATM/CHK2 pathway is primarily activated by double-strand breaks (DSBs), while the ATR/CHK1 pathway is triggered in response to replication fork collapse. Following DNA double-strand breaks (DSB), the ATM protein is activated by autophosphorylation, which then activates CHK2. The p53 tumour suppressor, a major effector of the DNA damage response (DDR) pathway, is expressed at low levels and in an inactive form during normal conditions. Both ATM and CHK2 phosphorylate p53, causing p53 protein stabilization and activation. Activated p53 arrests the cell cycle by inducing cell cycle inhibitors such as p21/CIP1. The DDR acting at the checkpoints normally allow the cell to repair its damaged DNA or alternatively undergo apoptosis [38].

## 3. Resistance to ionising irradiation in malignant and embryonic stem cells is associated with weak DNA damage checkpoints, reprogramming-senescence interplay, and reversible polyploidy

Whereas normal healthy somatic cells have the fate indicated above, the response of malignant cancer cells can differ leading to treatment resistance. Current data suggest that resistance can be induced in malignant tumour cells by reprogramming to an ESC-like state accompanied by WGD [14,21,39]. As illustrated in Fig. 1 A, B, the master regulator of embryonic stemness OCT4 can be induced in the Burkitt’s lymphoma mtTP53 cell line Namalwa alongside polyploidisation after 10-Gy irradiation [21]. In this study, there was up-regulated OCT4 in PML bodies alongside Nanog and SOX2 network, while all-trans-retinoic acid (an OCT4 antagonist) can disrupt Nanog nuclear localisation, and subsequently cell survival. Similarly, polyploidy-associated reprogramming induced by irradiation in breast cancer cells can be partially suppressed by Notch inhibition [14,39].

**Figure 1.**
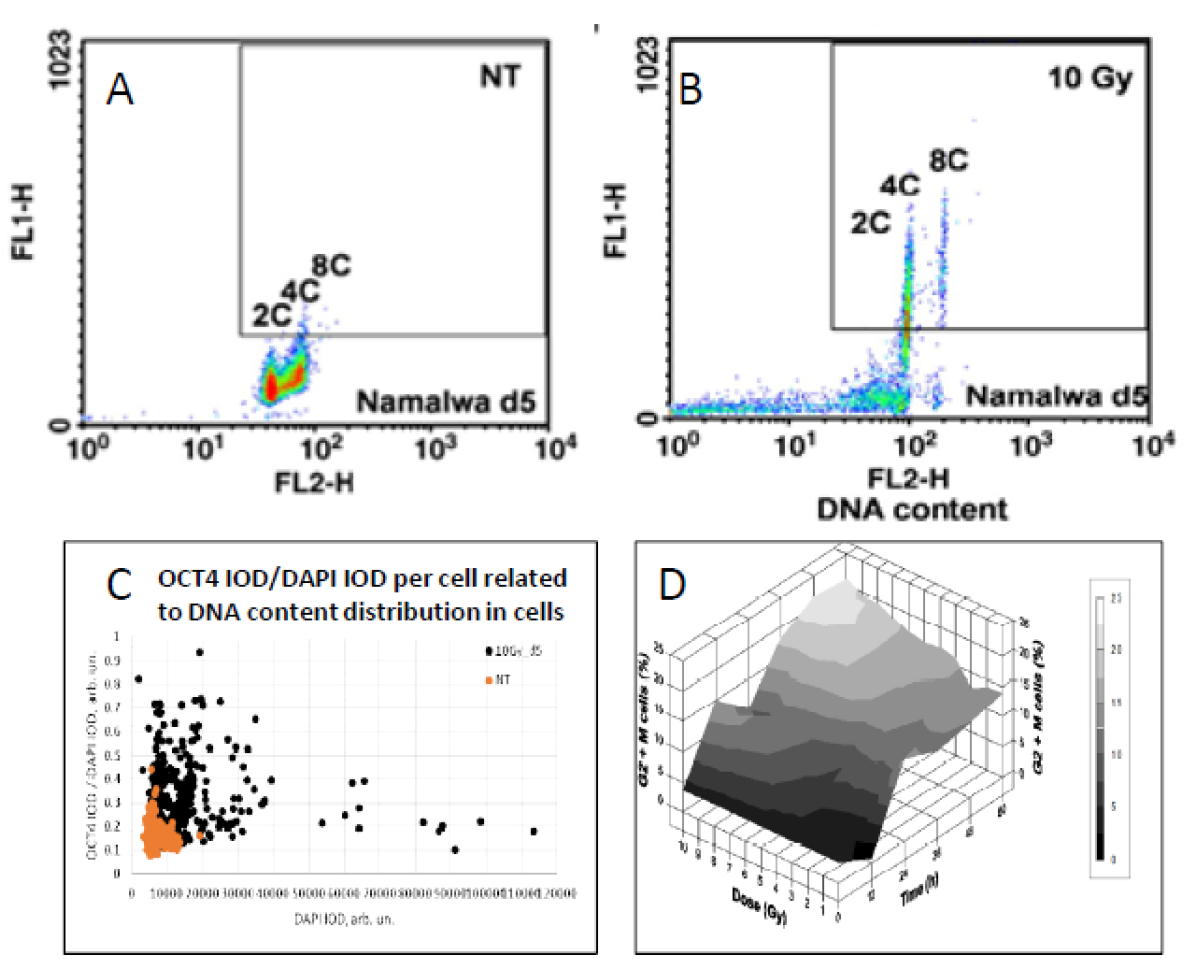
The similarity of responses to acute Irradiation (10 Gy, 2 Gy/min) in the malignant human Burkitt’s lymphoma cell line Namalwa and benign rat liver progenitor stem line WB-F344. Radiation-induced Oct4 upregulation in Namalwa cells as revealed by flow cytometry: panel **A** ‒ unirradiated cells (control); panel **B** ‒ irradiated cells on day 5 post-irradiation. According to the extent of FL1-signal (immunofluorescence from Oct4), Oct4 is predominantly expressed in polyploid 4C and 8C cells whose DNA content were determined by propidium iodide staining for DNA (FL2-signal) [from Salmina et al. 2010]. Panel **C**: radiation-induced Oct4 upregulation in WB-F344 cells as revealed by two-parametric image analysis of integrated optical densities (IOD): represented as Oct4 (IOD) / DAPI (IOD) versus DAPI (IOD). Panel **D**: radiation-induced G2/M delay in WB-F344 cells which is dose- and time-dependent [from Gerashchenko et al. 2004].

Strikingly similar post-irradiation effects were found in the rat liver epithelial cell line WB-F344, which is a hepatic tissue-specific progenitor capable of differentiating into hepatocytes and cholangiocytes [19]. This wtTP53 cell line, benign and incapable of inducing tumours *in* vivo, was shown to be radioresistant [40]. In common with genotoxic resistant cancers, the prominent feature of WB-F344 cells is a radiation dose-dependent enhancement of polyploidisation and micronucleation [40]; [19]. In this study, along with polyploidization, there was also up-regulation of the stemness transcription factors Oct4 and Nanog following 10-Gy irradiation [19], particularly in the polyploid fraction (Fig. 1C). Thus, while one is malignant (Namalwa), the other benign (WB-F344), these two radioresistant cell lines are capable of reprogramming – evoking induction of ESC-type stemness alongside polyploidy. Finally, there was radiation dose-dependent delay at the G2/M checkpoint (Fig. 2D) that preceded polyploidisation, and the same was found for malignant Burkitt’s lymphomas [13,20]. This response, characteristic for both resistant cell lines, is indicative of: (1) the weakness of the G1 checkpoint resulting in cell accumulation in G2M and, concurrently (2) the insufficiency of the G2M damage checkpoint, the main DSB sensor and actor, showing the tolerance to DNA DSBs and allowing transition to polyploidy. It is worth noting that persistently tolerable DDR signalling is a characteristic hallmark of ACS [26].

**Figure 2.**
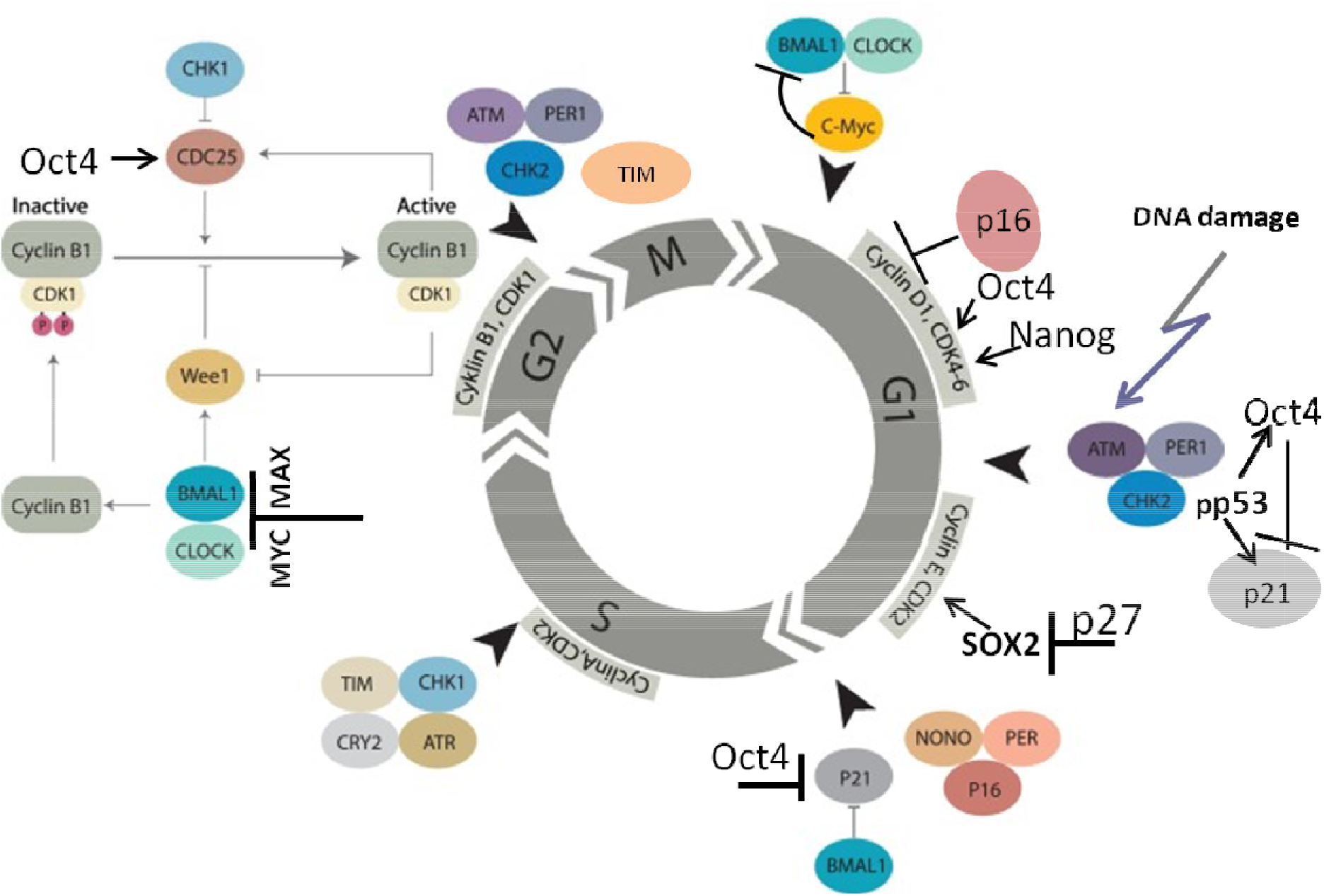
Molecular linkage between the regulators of the cell cycle in the embryonal (cancer) stem cells with the checkpoints adapted by basic stemness transcription factors in their relationship with CDK inhibitors (not all of them are shown) and Circadian Clock (adapted from [58] under Creative Common License). The details of the action of Circadian Clock regulators in DNA damage checkpoints and WGD are reviewed in sections 7 and 8.

Therefore, it is not surprising that our studies on the DDR, using three *in vitro* models of human lymphoma, ovarian embryonic carcinoma PA1 and rat liver stem cell lines [19,21,35], and *in vivo*, basal primary breast cancer (resistant to neoadjuvant therapy) [41], demonstrate reversible polyploidy with the interplay between Nanog and the senescence marker p16INK4A, and between Oct4 and p21CIP1 in the same or neighbouring cells, suggesting competitive oscillations between these opposing regulators of stemness and senescence [15].

Concluding this section, our studies on malignant and therapy-resistant Burkitt’s lymphoma *in vitro*, basal primary breast cancers, the ovarian embryonic carcinoma PA1, and the epithelial liver stem line WB344 revealed the consistent features – embryonic-type stemness (reprogramming) concurrent with senescence, attenuation of the DDR and transient polyploidisation. To better dissect this common mechanism, the regulation of the cell cycle checkpoints in ESCs will be reviewed in the next section.

## 4. Embryonic stem cells (ESCs) have defective cell cycle checkpoints that favour DNA damage tolerance and a shift to polyploidy

There is a body of evidence indicating that ESCs have a short G1-phase and weak or absent G1/S checkpoint, long S-phase, and weak intra-S and G2/M checkpoints [42–44]. In response to stress, ESCs have a tendency to undergo mitotic slippage from the spindle checkpoint, shifting to G1-tetraploidy at a specific stage with non-degradable cyclin B1, which protects ESCs from mitotic catastrophe [45]. In irradiated tumour cells, this stage is similar to endo-prometaphase [46]. Under stress, ESCs epigenetically switch off the p53 function [42,44]. The same is known for even TP53 wild-type tumours [47] whose response to DDR is very similar to the response of ESC. Both seemingly lack robust regulation of the cell cycle, tolerate DNA damage (thus display ACS) and when challenged attain polyploidy. Presumably, the inherent risk of genome instability that this brings is offset by and required for, their strategy for survival reliant upon explorative adaptation which demands the freedom of choice [15,48]. Induction of stemness in the damaged tumour cells in many ways is akin to the induced reprogramming by Yamanaka factors in normal cells [49] which also simultaneously causes DNA damage-tolerating senescence [26] that paradoxically is indispensable for its induction [28,29].

Transcription factors of the basic embryonal stemness network also possess the properties of cyclin-kinases or can otherwise overcome the senescence-driving and cell cycle-arresting cyclin-kinase inhibitors of the corresponding checkpoints. In particular, OCT4 induces the adaptation of the G1/S checkpoint by activating Cdk2 in the yclin E/Cdk2 complex [50] and enhancing the transcription of cyclin-kinases CDK4 and CDC25A [44,51]. OCT4 also toggles p21CIP1 ([52] in a p53-dependent (DDR-induced) manner [34,35,53]. Nanog activates Cdk6 by direct binding by ts C-domain [44,54], thus competing in the G1/S checkpoint with p16INK4a, which inhibits cyclin D. In the DDR, p16 is also activated by exaggerated expression of p21 and can cause terminal senescence [55]. Concurrently, together with IL-6, secreted by senescent cells, p16 is paradoxically indispensable for reprogramming [28]. In turn, SOX2 directly interacts with p27 (KIP1) in reprogramming to stimulate adaptation of the Cyclin E/Cdk2-dependent G1/S checkpoint [56] and also restricts the G2M checkpoint [57]. The most important activation of CDKs and opposing interactions between the embryonal stemness factors (OCT4, Nanog, and SOX2) with corresponding senescence regulators (p21, p16, p27) are shown in Fig.2.

This scheme will be used again in sections 7 and 8 describing the role of the CC in the cell cycle and WGD. The current analysis indicates that ESC cells tend to adapt to the checkpoints of the normal cell cycle, especially as part of their DDR.

“He who dares wins” (*qui audet vincit*). Mitotic slippage (MS) represents a transition compartment between the mitotic cell cycle and reversible polyploidy in the tumours undergoing DDR-mediated reprogramming. Three additional issues about MS need to be understood: (1) how the centrosomal cycle is affected? (2) what happens to the telomeres? (3) what is occurring with the biological time upturning from cell senescence for the birth of a new mitotic clone?

## 5. Mitotic slippage activates the cGAS-STING pathway and lifting the Hippo-surveillance of diploid mitosis favours polyploidisation

ASCs were shown to release heterochromatin particles into the cytoplasm inducing autophagic lysosome activity [59] and production of cytoplasmic DNA. This activates the cytosolic DNA-sensing innate immunity cGAS-STING pathway, producing diverse interferons and inflammatory cytokines [60][61]. The accompanying ACS-associated degradation of nuclear lamin B favours mitotic slippage and micronucleation of such cells, resetting interphase in a tetraploid state [62]. Interestingly, during mitotic slippage, the cGAS-STING-induced type I interferons reciprocally cooperate with the tumour suppressing Hippo pathway by activating LATS 1-2. In particular, the Aurora-A-Lats1/2-Aurora B axis pivotal for accurate coordination between chromosome segregation, karyo- and cytokinesis in anaphase, and midbody abscission in telophase becomes dysregulated, leading, also through the nuclear transition of YAP1, to bi-nuclearity, multinuclearity, and fusion of daughter nuclei [63–67]. The activated LATS1,2 instigate in addition the dysfunction of p53 [68] to promote cell migration.

## 6. Under-replication, erosion and recovery of ACS-compromised telomeres in mitotic slippage and transient polyploidy through transient alternative telomere lengthening

Cancer cell lines undergoing mitotic slippage accompanied by the cytoplasmic release of chromatin after genotoxic challenge also exhibit the under-replication of DNA in the late S-phase [35,69]. Under-replication of heterochromatin has been widely described in plants and insects, and Walter Nagl [70] indicated that it was always and only associated with the endocycle. Recent studies on Drosophila polyploid cells associate telomere under-replication with inhibition of replication fork progression and control of DNA copy number [71,72].

ACS was defined by Campisi [73] as cell stress that is characterised by compromised shortened telomeres which may be induced by oncogenic stress or DNA damage. As such, it appears that telomere erosion stemming from the heterochromatin under-replication may, in fact, result from replication stress reported in cancer development and treatment [74], occurring in the S-phase preceding polyploidisation by mitotic slippage (MS) in the same or rather (as observed) next cell cycle. Tam and colleagues [75] likely were the first to define ACS as a reversible process that is determined by the balance of biological molecules which directly or indirectly control telomere length and telomerase activity, by altering gene expression and/or modulating the epigenetic state of the chromatin. Our studies on the MDA-MB-231 breast cancer cell line treated with the Topoisomerase II inhibitor doxorubicin [69] revealed telomere ends enriched in DSBs massively casting off during MS together with the telomere capping protein TRF2 and the telomerase catalytic subunit TERT (Fig. 3 A-C). In the inter-and post-MS polyploid cells, restoration of the telomeres by alternative telomere lengthening (ALT) marked by specific TRF2-positive PML bodies was found (Fig.3D). It was followed by the recovery of TERT activity in the cells returning into the mitotic cell cycle (Fig.3 E, F). Importantly, in this interim process of telomere restoration through ALT-driven homologous recombination, the telomere ends of the chromosomes were found closed [69]. Telomere shortening in diploid somatic cells is associated with the linear chromosome end replication problem, cutting telomeres in each cell cycle by ~ 50 bp [76]. This process is the molecular basis underpinning the Hayflick limit [77], permitting cells to replicate only a limited number of times, proportional to the species lifespan. So, with the “trick” of under-replication signalling ACS and transient ALT, the chromosome end problem and the Hayflick (mortality) limit may be circumvented by polyploid tumour cells.

**Figure 3.**
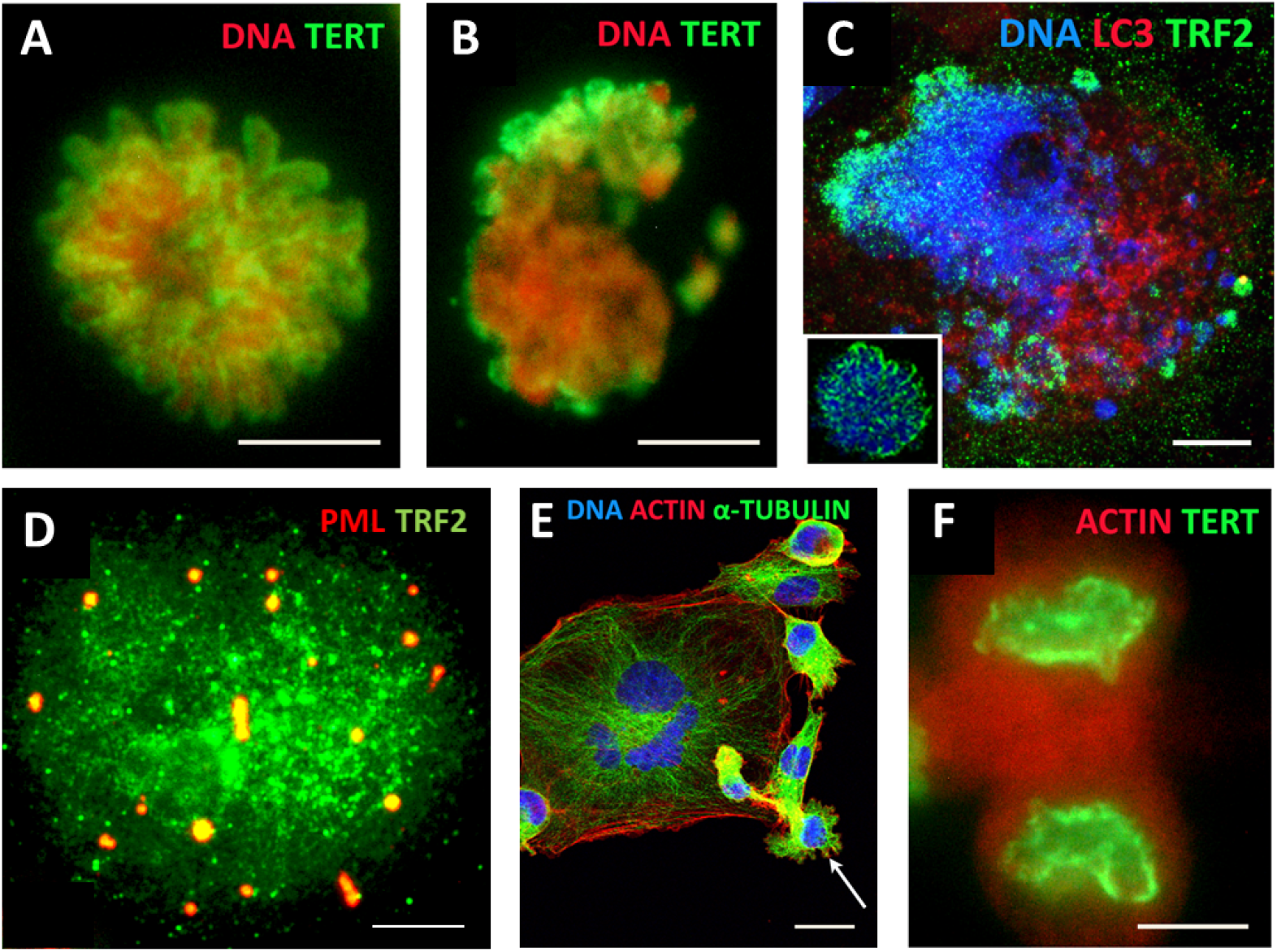
MDA-MB-231 breast cancer cell line (**A**) TERT-positive metaphase in control cells (DNA counterstained by propidium iodide); (**B**) mitotic slippage with poor TERT nuclear and enriched cytoplasmic DNA staining on Day 5 after DOX treatment; (**C**) preferential release of the telomere shelterin-TRF2-associated chromatin into the cytoplasm on Day 7 after DOX treatment (insert: normal metaphase); (**D**) polyploid cell, marked by specific TRF2-positive PML bodies showing restoration of the telomeres by alternative telomere lengthening (ALT); (**F**) TERT-positive escape telophase cell on Day 22 after DOX treatment; (**E**) A giant multinuclear cell is budding subcells (arrow). Bars: (A-D, F) = 10 μm; (E) = 25 μm.

A positive regulator of the telomere length Sirtuin 1, a NAD-dependent histone deacetylase (HDAC), directly binds to telomere repeats and attenuates telomere shortening associated with mouse ageing; this effect is dependent on telomerase activity [78]. At the same time, SIRT1 is very tightly associated with the regulation of the main cellular pacemaker - the CC. To analyse this aspect, we must first briefly describe the inner workings of this remarkable clock in the normal and ESC cell cycle (the latter is induced in tumours by DDR as described above in section 4).

## 7. The circadian clock (CC) paces the mitotic cell cycle, DDR checkpoints, and reciprocally, the TERT-dependent Hayflick limit count. It is absent in ESC and germ cells and likely becomes dis-engaged and then restored (by reversible polyploidy) in cancer cells

The bi-phasic CC is an autoregulatory transcriptional feedback loop-based oscillator involved in pacing the processes of living organisms with 24-hour rhythmicity [79]. The CC also regulates the cell cycle and couples various metabolic oscillations with shorter ultradian periodicity [80].

The core structure of the CC’s molecular oscillator contains a transcription activator arm, made up of BMAL1 and CLOCK, and a transcription repressor arm consisting of PER (Period) and CRY (Cryptochrome) genes. The heterodimeric complex of BMAL1 and CLOCK, which are basic helix-loop-helix transcription factors, binds the promoters and activates the expression of PER1, PER2, PER3, CRY1 and CRY2, that, in turn, heterodimerize into PER/CRY complexes, translocate into the nucleus and repress BMAL1/CLOCK [81]. The concentration of PER and CRY proteins is regulated by E3 ubiquitin ligases, resulting in their eventual depletion and BMAL/CLOCK1 reactivation [82]. A second, adjacent feedback loop involves nuclear receptors that bind DNA in a rhythmic manner - the activating (retinoic acid-dependent) RORs and repressive REV-ERBs [83]. These nuclear receptors regulate the expression of BMAL1 and NFIL3 and are themselves rhythmically regulated by the action of NFIL3, CLOCK, BMAL1 and DBP. In this way, the expression patterns of the clock components induce oscillatory behaviours in their downstream interactants [58,79]. The RORs-REV/ERBs loop as activated by retinoic acid is likely coupling CC with somatic cell differentiation. The CC in general is susceptible to stress conditions - the circadian cortisol-mediated entrainment of ultradian transcription pulses that provide the normal feedback regulation of cellular function is then lost [84].

Several genes of the CC deliver the strictly synchronized oscillation frequencies of the cell cycle [58,85] and participate in the regulation of the DNA damage checkpoints [80,86] as presented in Fig.2 [58]. The CC becomes dysfunctional in reprogramming induced by Yamanaka transcription factors [87]. Interestingly, circadian oscillation is also not detectable in ESCs until somite differentiation starts [88]. This may be related to the overexpressed ESC transcription factors speeding the cell cycle and forcing adaptation of its checkpoints as discussed in section 4 and illustrated in Fig.2. In addition, the direct competition of the main reprogramming transcription factor, MYC/MAX, with the CLOCK/BMAL1 dimer [80] in the G1/S and G2M checkpoints [89], which can be overcome through upregulated MYC [90](as designated on Fig.2), should be highlighted.

The loss of circadian rhythms impairs Hippo signalling, destabilizes p53 [91] and potentiates tumour initiation [92]. On the contrary, *in vitro* differentiation of ESCs induces cell-autonomous robust circadian oscillation [93]. It is important to note that besides ESC, the CC is also not functional in normal primordial germ cells (PGC) and both male and female gonocytes [94–96] suppressing somatic specification; the germline-specific protein PIWIL2 suppresses circadian rhythms [97] by inactivating the BMAL1 and CLOCK genes.

Noteworthy, in mammalian sperm, the telomere ends are joined, forming looped chromosomes [98], like those observed in mitotic slippage of cancer cells [69] and also in bi-parental bi-chromatid genome segregation (with disabled spindle) found by us alongside conventional mitoses in ovarian embryonal carcinoma [2]. Bi-parental gonomery was also described in glutamine-deprived normal fibroblasts initiating tetraploidy and suggested tumour conversion [99].

Interestingly, early mammalian embryos also display segregation of biparental genomes in the first short cleavage cycles [100] and also lack circadian regulation, which initiates in late embryos, tightly coupled to somatic cellular differentiation (in particular, somitogenesis) [101] or in vitro induced differentiation [87].

The above mentioned telomere-specific nicotinamide adenine dinucleotide (NADþ)-dependent HDAC SIRT 1, maintaining telomeres through telomerase activity, was found to interact with CLOCK and to be recruited to circadian promoters in a cyclic manner [102]. In particular, Wang RH et al. [103] showed that *Sirt1*-deficient mice exhibited profound premature ageing and enhanced acetylation of histone H4 in the promoter of *Per2*, the latter which leads to its overexpression; in turn, Per2 suppresses *Sirt1* transcription through binding to the *Sirt1* promoter at the Clock/Bmal1 site. This negative reciprocal relationship between SIRT1 and PER2 observed also in human hepatocytes, may perform the Hayflick limit count by CC.

We can subsequently rationalise that telomere shortening in ACS slows the circadian time-count and that further interruption of telomerase maintenance by TERT in MS substituted by recombination-based ALT with closed telomere ends should interrupt CC (arresting the biological time pace) while returning to the TERT mechanism in depolyploidized offspring restoring the mitotic cycle [69] should resume the CC oscillation and hence the Hayflick limit count. This manipulation of biological time in MS is reminiscent of a “death loop” in aviation.

## 8. The circadian clock in the context of mammalian polyploidy and cancer

### 8.1. The reciprocal regulation of polyploidy and CC activity in non-malignant tissues

The competitive antagonism of the overexpressed stemness/reprogramming master factor dimer MYC/MAX with CLOCK/BMAL1, which is the core component of the CC’s activation arm and a regulator of the G2M DNA damage checkpoint, is likely to play a key role in impairing the CC in stem cells (including stressed cancer cells that have undergone reprogramming), where stemness features were shown to be tightly coupled to deregulation of the cell that leads to polyploidy. The Timeless (TIM) gene was shown to be involved in the S-phase checkpoint [104]. The circadian clock proteins PER1, PER2 and PER3 are involved in the ploidy regulation of non-cancerous liver cells, and their inactivation results in rampant polyploidization (both in terms of polyploidization frequency and increased ploidy counts in the polyploid hepatocytes) [105]. It is also important to mention that of the 16 core genes of the circadian clock (CLOCK, ARNTL (BMAL1), ARNTL2, NPAS2, NR1D1, NR1D2, CRY1, CRY2, DBP, TEF, RORA, RORB, RORC, PER1, PER2, and PER3) 50% can be found in the list of bivalent genes [106] allowing rapid cell fate change. Interestingly, polyploidy (the endocycle) in plants was shown to decelerate the circadian rhythm [107]. In addition, evidence from mouse and human transcriptome analyses, suggests that the deregulation of the circadian clock promotes polyploidization and *vice versa* [105,108]. In turn, polyploidy in normal tissues like the mammalian heart and liver is associated with up-regulated c-Myc and the stemness and cancer-linked EMT targets [109]. The role of the CC in cell cycle integrity and DDR signalling is further showcased by its involvement in DNA repair after ionizing irradiation damage (by inducing DDR-signaling genes) (Fig.2) [58,110,111]).

The CC is notably deregulated in cancer [112,113] and mediated by Ras-oncogene (and mediated by Ras-oncogene [114]. In turn, perturbation of the CC is in itself carcinogenic [115,116]. Meta-analysis of 7476 cancer cases from 36 sources [117] revealed that low expression of PER1 and PER2 correlates with poor differentiation, worse TNM stage, metastases, and reduced patient survival.

Overall, the currently available information on the connection between the CC, stemness and the cell cycle, as well CC deregulation in cancer, leads us to suggest that circadian deregulation in human cancer may be largely associated with its polyploidy component as it is in normal mammalian heart and liver. In the next section, we describe an attempt to investigate this hypothesis through bioinformatics analysis of primary cancers.

### 8.2. Circadian deregulation correlates with polyploidization (whole-genome doubling) in malignant tumour patient samples

In order to investigate the possible connection between polyploidy and CC deregulation in cancer, it was first necessary to calculate the measure of circadian deregulation. To that end, we used the Cancer Genome Atlas (TCGA), a large-scale collection of omics and clinical data on over 30 types of malignancies from over 11000 patients [118]. TPM-normalized Rsubread-processed TCGA gene expression data was obtained from the GSE62944 GEO dataset [119]. In order to ensure statistical power, only TCGA transcriptomic datasets counterpart by at least 35 available normal samples were selected, resulting in a final cohort of 11 cancer types and 6667 samples (613 normal and 6054 tumours).

Circadian deregulation in TCGA cancer samples was determined using the CCD method and deltaccd R package developed by Shilts et al. [112], which compares core CC gene co-expression (Spearman rank-based correlation) between samples used in the study and a pan-tissue reference matrix calculated from 8 normal mouse datasets with available time data. The Euclidean distance between CC gene correlation vectors of the samples and the mouse reference is referred to as the Clock Correlation Distance (CCD). The difference between normal VS reference CCD, and the tumour VS reference CCD, known as the CCD, serves as a coefficient of circadian dysregulation, with the “difference of differences” approach effectively negating the nuance of mouse-human comparison, and accepting the common regulation of CC in mammals [120].

Tumour ploidy calculated from copy-number data using the ABSOLUTE algorithm [121] was obtained from [122] and the relationship between the values of scaled CCD for each of the 11 tumour types and the respective proportion of samples with at least one WGD was investigated using Spearman correlation analysis.

The results revealed a statistically significant positive correlation (Spearman’s rho=0.83; p<0.01) between WGD and CC deregulation (Fig.4). While correlation does not necessarily equal causation, such a result seems logically sound when taking into account the known associations between polyploidy and the CC in normal tissues, deregulation of CC in cancers, as well as the impact of polyploidy on cancer evolution.

**Figure 4.**
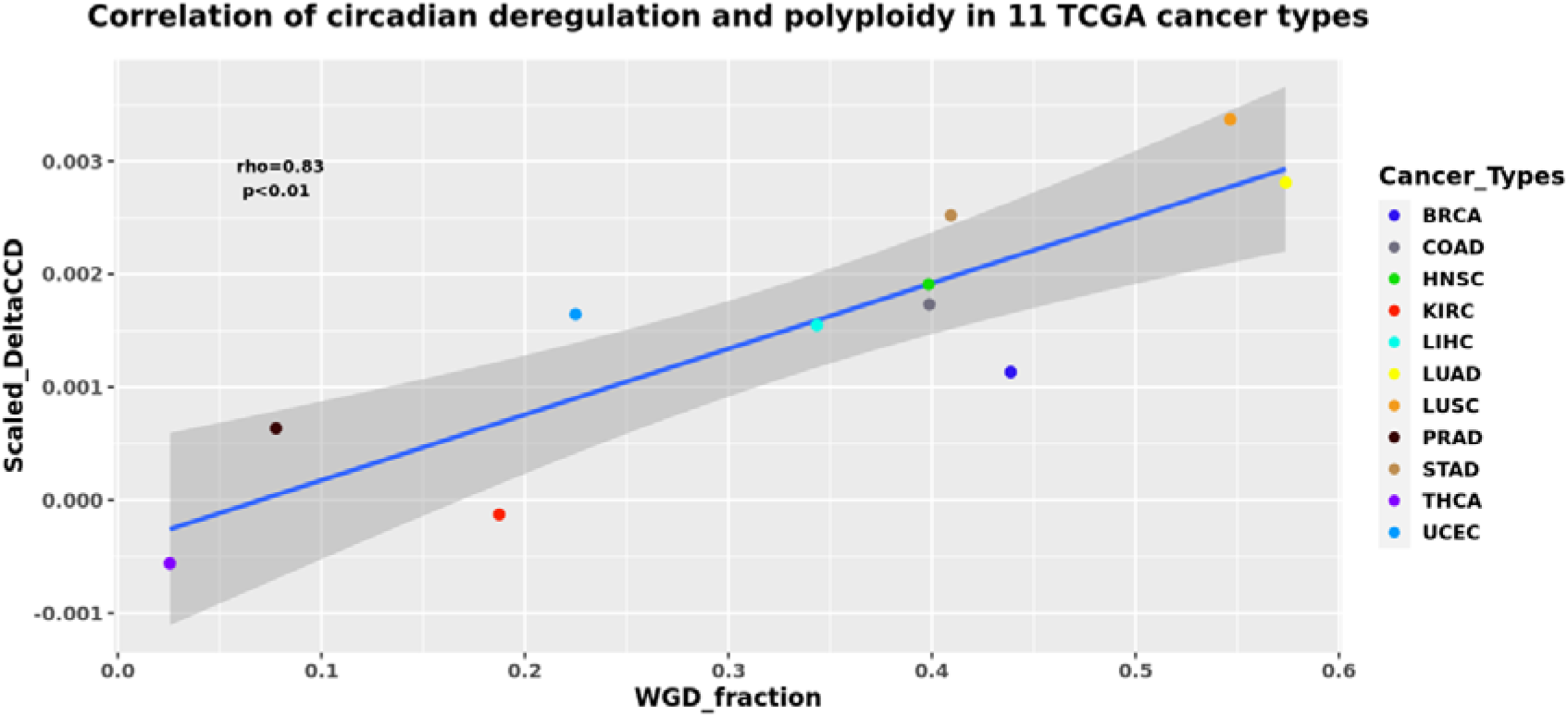
The CCD coefficient of circadian deregulation positively correlates with the proportion of WGD in the samples of 11 tumour types from The Cancer Genome Atlas (TCGA) database. BRCA- breast carcinoma, COAD- colon adenocarcinoma, HNSC - head and neck squamous cell carcinoma, KIRC - kidney renal cell carcinoma, LIHC - liver hepatocellular carcinoma, LUAD - lung adenocarcinoma, LUSC - lung squamous cell carcinoma, PRAD - prostate adenocarcinoma, STAD - gastric adenocarcinoma, THCA - thyroid carcinoma, UCEC - uterine corpus endometrial carcinoma.

It is interesting, as mentioned above, that the non-functional CC also characterises the features of germ cells and early embryos - the beginnings of a life cycle. This reintroduces the legacy of the oldest embryonal theory of cancer and its parthenogenetic variants [22–24,123–126]. Furthermore, the connection between the senescence and mitotic slippage-induced cGAS-STING-Type I Interferon pathway and one of its targets - the transmembrane gene family Fragilis (IFITM3) that is involved in the early commitment of PGCs [127,128] and expressed in oogonial stem cells [129] may be added as further evidence of soma-to-germ transition in cancer.

## 9. Conclusion

Currently, data regarding the role and importance of the de-regulation of the CC in cancer are accumulating. Its possible role in treatment resistance was analyzed here. We evaluated the reprogramming (stemness) of genotoxically treated cancer cells with the impact on the cell cycle - overcoming the G1/S checkpoint, adapting DNA damage checkpoints and tolerating DNA damage, thus outlining the features of ASC with eroded telomeres, which pushes cells through MS into polyploidy. Next, we evaluated this observation with a correlation between cancer polyploidy and deregulation of CC using bioinformatics analysis of the TCGA database of primary cancers. More lethal cancers have the highest WGD proportion, correlating with the largest deregulation of the CC. The data shows that to get there after receiving genotoxic stress a cancer cell should perform a “death loop” - falling out of the canonical mitotic cell cycle, (normally driven by the CC), into a polyploidy cycle with decelerated or non-functional CC followed by a return to the mitotic cycle. In some way, this “fall” resets the cell to a ‘timeless state', the likes of which are normally displayed only by germ cells and early embryos which exhibit zero CC oscillation. Return to the biological time pace, which is normally associated with counting of the Hayflick limit, needs the telomeres (having undergone attrition by ACS) to be restored and again coupled to TERT. This telomere restitution mechanism may be driven by ALT coupled to a kind of meiotic homology search and recombination [69]. This need can also explain the rich germline signature and ectopic expression of meiotic genes in cancer correlating with both polyploidy and poor prognosis [130,131], and also corresponding to the blastomere-like features of polyploid giant cancer cells [16,22,23].

ALT combined with the meiotic type recombination (possibly, inverted meiosis [69] can restrict aneuploidy and return the depolyploidized offspring to the Hayflick limit counter, onto the CC oscillations again. Within this logic, the various requirements of the above-discussed mechanisms of resistance to anticancer treatments (WGD, senescence, reprogramming, deregulated CC, attrition and recovery of telomeres) may be provisionally met (Fig.5, Graphical Abstract). Hopefully, this analysis will serve to inspire further assessments of these inter-relationships in cancer research and lead to better strategies of cancer prevention, prognosis, and treatment.

**Fig.5.**
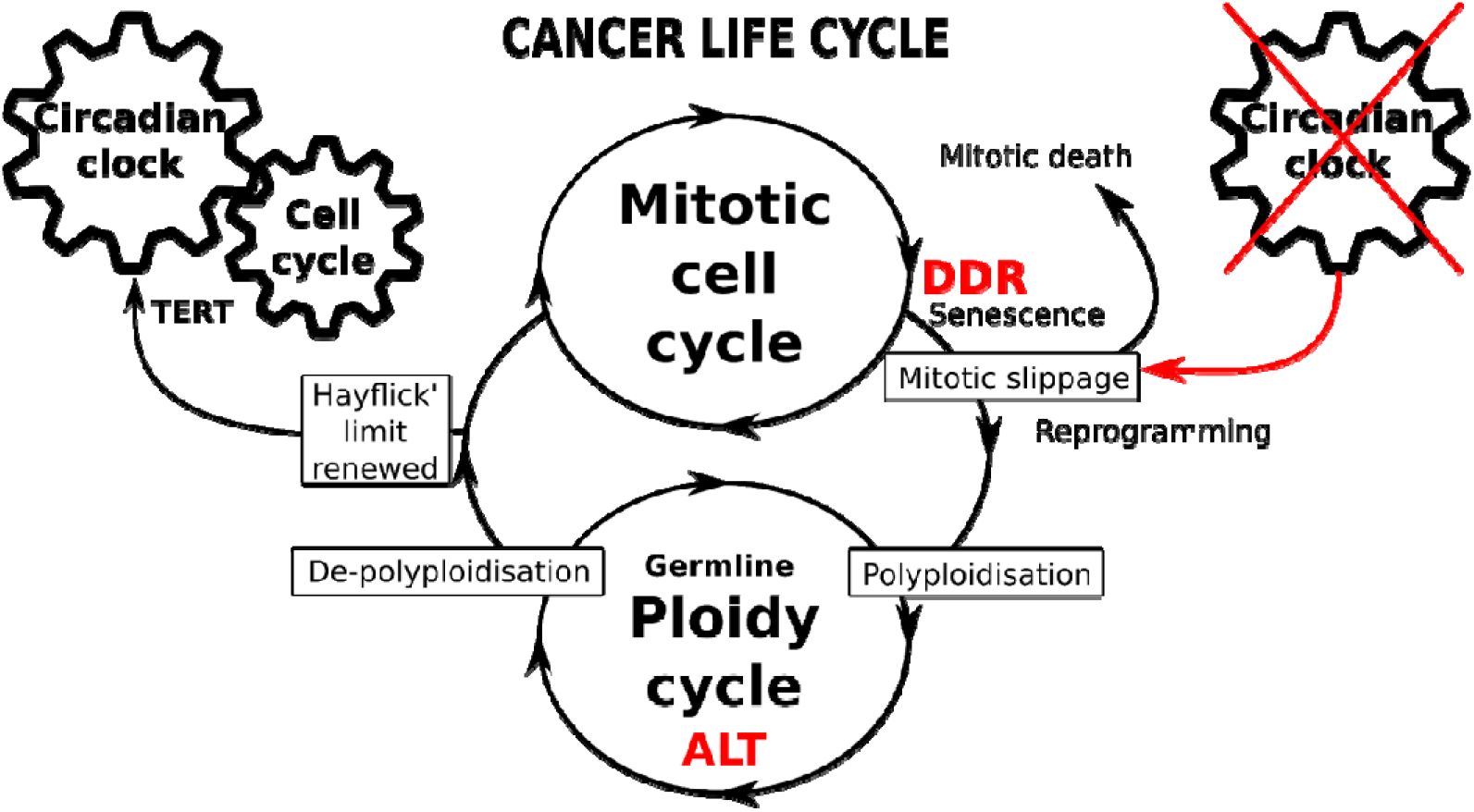
Schematic of the immortal cancer life-cycle composed of two reciprocally joined mitotic and ploidy cycles. The mitotic cell cycle is driven by the circadian clock, in particular operating the telomerase (TERT-dependent) telomere maintenance pathway. The transition from mitotic to ploidy cycle is occurring after adapted DNA checkpoints of DNA damage response (DDR), through mitotic slippage coupling accelerated cell senescence (with compromised telomeres) and reprogramming with whole-genome duplications. Transition into the ploidy cycle, featured by germline expression signature, is associated with interruption of circadian clock and restoration of eroded telomeres by alternative telomere lengthening (ALT). Return of depolyploidised offspring to mitotic cycle restores the TERT-pathway and the CC-driven count of Hayflick limit.

## Acknowledgements

Jēkabs Krīgerts is acknowledged for the cytometry of WB344 cells.

## References

1. Erenpreisa, J.; Cragg, M.S. Three Steps to the Immortality of Cancer Cells: Senescence, Polyploidy and Self-Renewal. Cancer Cell Int. 2013, 13, 92.

2. Salmina, K.; Huna, A.; Kalejs, M.; Pjanova, D.; Scherthan, H.; Cragg, M.S.; Erenpreisa, J. The Cancer Aneuploidy Paradox: In the Light of Evolution. Genes 2019, 10, doi:10.3390/genes10020083.

3. Bielski, C.M.; Zehir, A.; Penson, A.V.; Donoghue, M.T.A.; Chatila, W.; Armenia, J.; Chang, M.T.; Schram, A.M.; Jonsson, P.; Bandlamudi, C.; et al. Genome Doubling Shapes the Evolution and Prognosis of Advanced Cancers. Nat. Genet. 2018, 50, 1189–1195.

4. Lee, H.O.; Davidson, J.M.; Duronio, R.J. Endoreplication: Polyploidy with Purpose. Genes Dev. 2009, 23, 2461–2477.

5. Van de Peer, Y.; Mizrachi, E.; Marchal, K. The Evolutionary Significance of Polyploidy. Nat. Rev. Genet. 2017, 18, 411–424.

6. Pienta, K.J.; Hammarlund, E.U.; Axelrod, R.; Brown, J.S.; Amend, S.R. Poly-Aneuploid Cancer Cells Promote Evolvability, Generating Lethal Cancer. Evol. Appl. 2020, 13, 1626–1634.

7. Zhang, S.; Mercado-Uribe, I.; Xing, Z.; Sun, B.; Kuang, J.; Liu, J. Generation of Cancer Stem-like Cells through the Formation of Polyploid Giant Cancer Cells. Oncogene 2014, 33, 116–128.

8. Mirzayans, R.; Andrais, B.; Murray, D. Roles of Polyploid/Multinucleated Giant Cancer Cells in Metastasis and Disease Relapse Following Anticancer Treatment. Cancers 2018, 10, doi:10.3390/cancers10040118.

9. Díaz-Carballo, D.; Saka, S.; Klein, J.; Rennkamp, T.; Acikelli, A.H.; Malak, S.; Jastrow, H.; Wennemuth, G.; Tempfer, C.; Schmitz, I.; et al. A Distinct Oncogenerative Multinucleated Cancer Cell Serves as a Source of Stemness and Tumor Heterogeneity. Cancer Res. 2018, 78, 2318–2331.

10. Díaz-Carballo, D.; Gustmann, S.; Jastrow, H.; Acikelli, A.H.; Dammann, P.; Klein, J.; Dembinski, U.; Bardenheuer, W.; Malak, S.; Araúzo-Bravo, M.J.; et al. Atypical Cell Populations Associated with Acquired Resistance to Cytostatics and Cancer Stem Cell Features: The Role of Mitochondria in Nuclear Encapsulation. DNA Cell Biol. 2014, 33, 749–774.

11. Sundaram, M.; Guernsey, D.L.; Rajaraman, M.M.; Rajaraman, R. Neosis: A Novel Type of Cell Division in Cancer. Cancer Biol. Ther. 2004, 3, 207–218.

12. Illidge, T.M.; Cragg, M.S.; Fringes, B.; Olive, P.; Erenpreisa, J.A. Polyploid Giant Cells Provide a Survival Mechanism for p53 Mutant Cells after DNA Damage. Cell Biol. Int. 2000, 24, 621–633.

13. Erenpreisa, J.A.; Cragg, M.S.; Fringes, B.; Sharakhov, I.; Illidge, T.M. Release of Mitotic Descendants by Giant Cells from Irradiated Burkitt’s Lymphoma Cell Line. Cell Biol. Int. 2000, 24, 635–648.

14. Lagadec, C.; Vlashi, E.; Della Donna, L.; Dekmezian, C.; Pajonk, F. Radiation-Induced Reprogramming of Breast Cancer Cells. Stem Cells 2012, 30, 833–844.

15. Erenpreisa, J.; Salmina, K.; Anatskaya, O.; Cragg, M.S. Paradoxes of Cancer: Survival at the Brink. Semin. Cancer Biol. 2020, doi:10.1016/j.semcancer.2020.12.009.

16. Erenpreisa, J.; Cragg, M.S. Cancer: A Matter of Life Cycle? Cell Biol. Int. 2007, 31, 1507–1510.

17. Erenpreisa, J.; Cragg, M.S. MOS, Aneuploidy and the Ploidy Cycle of Cancer Cells. Oncogene 2010, 29, 5447–5451.

18. Edgar, B.A.; Orr-Weaver, T.L. Endoreplication Cell Cycles. Cell 2001, 105, 297–306.

19. Gerashchenko, B.I.; Salmina, K.; Krigerts, J.; Erenpreisa, J.; Babsky, A.M. INDUCED POLYPLOIDY AND SORTING OF DAMAGED DNA BY MICRONUCLEATION IN RADIORESISTANT RAT LIVER EPITHELIAL STEM-LIKE CELLS EXPOSED TO X-RAYS. Probl Radiac Med Radiobiol 2019, 24, 220–234.

20. Ivanov, A.; Cragg, M.S.; Erenpreisa, J.; Emzinsh, D.; Lukman, H.; Illidge, T.M. Endopolyploid Cells Produced after Severe Genotoxic Damage Have the Potential to Repair DNA Double Strand Breaks. J. Cell Sci. 2003, 116, 4095–4106.

21. Salmina, K.; Jankevics, E.; Huna, A.; Perminov, D.; Radovica, I.; Klymenko, T.; Ivanov, A.; Jascenko, E.; Scherthan, H.; Cragg, M.; et al. Up-Regulation of the Embryonic Self-Renewal Network through Reversible Polyploidy in Irradiated p53- Mutant Tumour Cells. Exp. Cell Res. 2010, 316, 2099–2112.

22. Erenpreisa, J.; Salmina, K.; Huna, A.; Jackson, T.R.; Vazquez-Martin, A.; Cragg, M.S. The “Virgin Birth”, Polyploidy, and the Origin of Cancer. Oncoscience 2015, 2, 3–14.

23. Niu, N.; Mercado-Uribe, I.; Liu, J. Dedifferentiation into Blastomere-like Cancer Stem Cells via Formation of Polyploid Giant Cancer Cells. Oncogene 2017, 36, 4887–4900.

24. Liu, J. The “life Code”: A Theory That Unifies the Human Life Cycle and the Origin of Human Tumors. Seminars in Cancer Biology 2020, 60, 380–397.

25. Roninson, I.B.; Broude, E.V.; Chang, B.-D. If Not Apoptosis, Then What? Treatment-Induced Senescence and Mitotic Catastrophe in Tumor Cells. Drug Resistance Updates 2001, 4, 303–313.

26. Campisi, J.; d’Adda di Fagagna, F. Cellular Senescence: When Bad Things Happen to Good Cells. Nat. Rev. Mol. Cell Biol. 2007, 8, 729–740.

27. Davoli, T.; de Lange, T. Telomere-Driven Tetraploidization Occurs in Human Cells Undergoing Crisis and Promotes Transformation of Mouse Cells. Cancer Cell 2012, 21, 765–776.

28. Mosteiro, L.; Pantoja, C.; de Martino, A.; Serrano, M. Senescence Promotes in Vivo Reprogramming through p16 and IL-6. Aging Cell 2018, 17, doi:10.1111/acel.12711.

29. Lee, S.; Schmitt, C.A. The Dynamic Nature of Senescence in Cancer. Nat. Cell Biol. 2019, 21, 94–101.

30. Moein, S.; Adibi, R.; da Silva Meirelles, L.; Nardi, N.B.; Gheisari, Y. Cancer Regeneration: Polyploid Cells Are the Key Drivers of Tumor Progression. Biochim. Biophys. Acta Rev. Cancer 2020, 1874, 188408.

31. Mosieniak, G.; Sliwinska, M.A.; Alster, O.; Strzeszewska, A.; Sunderland, P.; Piechota, M.; Was, H.; Sikora, E. Polyploidy Formation in Doxorubicin-Treated Cancer Cells Can Favor Escape from Senescence. Neoplasia 2015, 17, 882–893.

32. Mosieniak, G.; Sikora, E. Polyploidy: The Link between Senescence and Cancer. Curr. Pharm. Des. 2010, 16, 734–740.

33. Puig, P.-E.; Guilly, M.-N.; Bouchot, A.; Droin, N.; Cathelin, D.; Bouyer, F.; Favier, L.; Ghiringhelli, F.; Kroemer, G.; Solary, E.; et al. Tumor Cells Can Escape DNA-Damaging Cisplatin through DNA Endoreduplication and Reversible Polyploidy. Cell Biol. Int. 2008, 32, 1031–1043.

34. Jackson, T.R.; Salmina, K.; Huna, A.; Inashkina, I.; Jankevics, E.; Riekstina, U.; Kalnina, Z.; Ivanov, A.; Townsend, P.A.; Cragg, M.S.; et al. DNA Damage Causes TP53-Dependent Coupling of Self-Renewal and Senescence Pathways in Embryonal Carcinoma Cells. Cell Cycle 2013, 12, 430–441.

35. Huna, A.; Salmina, K.; Erenpreisa, J.; Vazquez-Martin, A.; Krigerts, J.; Inashkina, I.; Gerashchenko, B.I.; Townsend, P.A.; Cragg, M.S.; Jackson, T.R. Role of Stress-Activated OCT4A in the Cell Fate Decisions of Embryonal Carcinoma Cells Treated with Etoposide. Cell Cycle 2015, 14, 2969–2984.

36. Suzuki, M.; Boothman, D.A. Stress-Induced Premature Senescence (SIPS)--Influence of SIPS on Radiotherapy. J. Radiat. Res. 2008, 49, 105–112.

37. Weihua, Z.; Lin, Q.; Ramoth, A.J.; Fan, D.; Fidler, I.J. Formation of Solid Tumors by a Single Multinucleated Cancer Cell. Cancer 2011, 117, 4092–4099.

38. Barnum, K.J.; O’Connell, M.J. Cell Cycle Regulation by Checkpoints. Methods Mol. Biol. 2014, 1170, 29–40.

39. Vlashi, E.; Pajonk, F. Cancer Stem Cells, Cancer Cell Plasticity and Radiation Therapy. Semin. Cancer Biol. 2015, 31, 28–35.

40. Gerashchenko, B.I.; Azzam, E.I.; Howell, R.W. Characterization of Cell-Cycle Progression and Growth of WB-F344 Normal Rat Liver Epithelial Cells Following Gamma-Ray Exposure. Cytometry A 2004, 61, 134–141.

41. Gerashchenko, B.I.; Salmina, K.; Eglitis, J.; Huna, A.; Grjunberga, V.; Erenpreisa, J. Disentangling the Aneuploidy and Senescence Paradoxes: A Study of Triploid Breast Cancers Non-Responsive to Neoadjuvant Therapy. Histochem. Cell Biol. 2016, 145, 497–508.

42. Suvorova, I.I.; Grigorash, B.B.; Chuykin, I.A.; Pospelova, T.V.; Pospelov, V.A. G1 Checkpoint Is Compromised in Mouse ESCs due to Functional Uncoupling of p53-p21Waf1 Signaling. Cell Cycle 2016, 15, 52–63.

43. Liu, L.; Michowski, W.; Kolodziejczyk, A.; Sicinski, P. The Cell Cycle in Stem Cell Proliferation, Pluripotency and Differentiation. Nat. Cell Biol. 2019, 21, 1060–1067.

44. Neganova, I.; Lako, M. G1 to S Phase Cell Cycle Transition in Somatic and Embryonic Stem Cells. Journal of Anatomy 2008, 213, 30–44.

45. Mantel, C.; Guo, Y.; Lee, M.R.; Han, M.K.; Rhorabough, S.; Kim, K.S.; Broxmeyer, H.E. Cells Enter a Unique Intermediate 4N Stage, Not 4N-G1, after Aborted Mitosis. Cell Cycle 2008, 7, 484–492.

46. Erenpreisa, J.; Kalejs, M.; Cragg, M.S. Mitotic Catastrophe and Endomitosis in Tumour Cells: An Evolutionary Key to a Molecular Solution. Cell Biol. Int. 2005, 29, 1012–1018.

47. Kastan, M.B. Wild-Type p53: Tumors Can’t Stand It. Cell 2007, 128, 837–840.

48. Huang, S. Reprogramming Cell Fates: Reconciling Rarity with Robustness. Bioessays 2009, 31, 546–560.

49. Takahashi, K.; Yamanaka, S. Induction of Pluripotent Stem Cells from Mouse Embryonic and Adult Fibroblast Cultures by Defined Factors. Cell 2006, 126, 663–676.

50. Neganova, I.; Vilella, F.; Atkinson, S.P.; Lloret, M.; Passos, J.F.; von Zglinicki, T.; O’Connor, J.-E.; Burks, D.; Jones, R.; Armstrong, L.; et al. An Important Role for CDK2 in G1 to S Checkpoint Activation and DNA Damage Response in Human Embryonic Stem Cells. STEM CELLS 2011, 29, 651–659.

51. Greco, S.J.; Liu, K.; Rameshwar, P. Functional Similarities among Genes Regulated by OCT4 in Human Mesenchymal and Embryonic Stem Cells. Stem Cells 2007, 25, 3143–3154.

52. Keyes, W.M.; Wu, Y.; Vogel, H.; Guo, X.; Lowe, S.W.; Mills, A.A. p63 Deficiency Activates a Program of Cellular Senescence and Leads to Accelerated Aging. Genes Dev. 2005, 19, 1986–1999.

53. Baryshev, M.; Inashkina, I.; Salmina, K.; Huna, A.; Jackson, T.R.; Erenpreisa, J. DNA Methylation of the Oct4A Enhancers in Embryonal Carcinoma Cells after Etoposide Treatment Is Associated with Alternative Splicing and Altered Pluripotency in Reversibly Senescent Cells. Cell Cycle 2018, 17, 362–366.

54. Zhang, X.; Neganova, I.; Przyborski, S.; Yang, C.; Cooke, M.; Atkinson, S.P.; Anyfantis, G.; Fenyk, S.; Keith, W.N.; Hoare, S.F.; et al. A Role for NANOG in G1 to S Transition in Human Embryonic Stem Cells through Direct Binding of CDK6 and CDC25A. J. Cell Biol. 2009, 184, 67–82.

55. Kulaberoglu, Y.; Gundogdu, R.; Hergovich, A. The Role of p53/p21/p16 in DNA-Damage Signaling and DNA Repair. In Genome Stability; Elsevier, 2016; pp. 243–256 ISBN 9780128033098.

56. Li, H.; Collado, M.; Villasante, A.; Matheu, A.; Lynch, C.J.; Cañamero, M.; Rizzoti, K.; Carneiro, C.; Martínez, G.; Vidal, A.; et al. p27(Kip1) Directly Represses Sox2 during Embryonic Stem Cell Differentiation. Cell Stem Cell 2012, 11, 845–852.

57. She, S.; Wei, Q.; Kang, B.; Wang, Y.-J. Cell Cycle and Pluripotency: Convergence on Octamer-binding Transcription Factor 4 (Review). Mol. Med. Rep. 2017, 16, 6459–6466.

58. Farshadi, E.; van der Horst, G.T.J.; Chaves, I. Molecular Links between the Circadian Clock and the Cell Cycle. J. Mol. Biol. 2020, 432, 3515–3524.

59. Ivanov, A.; Pawlikowski, J.; Manoharan, I.; van Tuyn, J.; Nelson, D.M.; Rai, T.S.; Shah, P.P.; Hewitt, G.; Korolchuk, V.I.; Passos, J.F.; et al. Lysosome-Mediated Processing of Chromatin in Senescence. J. Cell Biol. 2013, 202, 129–143.

60. Dou, Z.; Ghosh, K.; Vizioli, M.G.; Zhu, J.; Sen, P.; Wangensteen, K.J.; Simithy, J.; Lan, Y.; Lin, Y.; Zhou, Z.; et al. Cytoplasmic Chromatin Triggers Inflammation in Senescence and Cancer. Nature 2017, 550, 402–406.

61. Kwon, J.; Bakhoum, S.F. The Cytosolic DNA-Sensing cGAS-STING Pathway in Cancer. Cancer Discov. 2020, 10, 26–39.

62. Lezaja, A.; Altmeyer, M. Dealing with DNA Lesions: When One Cell Cycle Is Not Enough. Curr. Opin. Cell Biol. 2021, 70, 27–36.

63. Steigemann, P.; Wurzenberger, C.; Schmitz, M.H.A.; Held, M.; Guizetti, J.; Maar, S.; Gerlich, D.W. Aurora B-Mediated Abscission Checkpoint Protects against Tetraploidization. Cell 2009, 136, 473–484.

64. Bui, D.A.; Lee, W.; White, A.E.; Harper, J.W.; Schackmann, R.C.J.; Overholtzer, M.; Selfors, L.M.; Brugge, J.S. Cytokinesis Involves a Nontranscriptional Function of the Hippo Pathway Effector YAP. Sci. Signal. 2016, 9, ra23.

65. Yabuta, N.; Mukai, S.; Okada, N.; Aylon, Y.; Nojima, H. The Tumor Suppressor Lats2 Is Pivotal in Aurora A and Aurora B Signaling during Mitosis. Cell Cycle 2011, 10, 2724–2736.

66. Vitale, I.; Galluzzi, L.; Castedo, M.; Kroemer, G. Mitotic Catastrophe: A Mechanism for Avoiding Genomic Instability. Nat. Rev. Mol. Cell Biol. 2011, 12, 385–392.

67. Ganem, N.J.; Cornils, H.; Chiu, S.-Y.; O’Rourke, K.P.; Arnaud, J.; Yimlamai, D.; Théry, M.; Camargo, F.D.; Pellman, D. Cytokinesis Failure Triggers Hippo Tumor Suppressor Pathway Activation. Cell 2014, 158, 833–848.

68. Furth, N.; Bossel Ben-Moshe, N.; Pozniak, Y.; Porat, Z.; Geiger, T.; Domany, E.; Aylon, Y.; Oren, M. Down-Regulation of LATS Kinases Alters p53 to Promote Cell Migration. Genes Dev. 2015, 29, 2325–2330.

69. Salmina, K.; Bojko, A.; Inashkina, I.; Staniak, K.; Dudkowska, M.; Podlesniy, P.; Rumnieks, F.; Vainshelbaum, N.M.; Pjanova, D.; Sikora, E.; et al. “Mitotic Slippage” and Extranuclear DNA in Cancer Chemoresistance: A Focus on Telomeres. Int. J. Mol. Sci. 2020, 21, doi:10.3390/ijms21082779.

70. Nagl, W. Endopolyploidy and Polyteny in Differentiation and Evolution: Towards an Understanding of Quantitative and Qualitative Variation of Nuclear DNA in Ontogeny and Phylogeny; North-Holland, 1978;.

71. Munden, A.; Rong, Z.; Sun, A.; Gangula, R.; Mallal, S.; Nordman, J.T. Rif1 Inhibits Replication Fork Progression and Controls DNA Copy Number in Drosophila. Elife 2018, 7, doi:10.7554/eLife.39140.

72. Das, S.; Caballero, M.; Kolesnikova, T.; Zhimulev, I.; Koren, A.; Nordman, J. Replication Timing Analysis in Polyploid Cells Reveals Rif1 Uses Multiple Mechanisms to Promote Underreplication in Drosophila. Genetics 2021, 219, doi:10.1093/genetics/iyab147.

73. Campisi, J. Aging, Cellular Senescence, and Cancer. Annu. Rev. Physiol. 2013, 75, 685–705.

74. Bartek, J.; Mistrik, M.; Bartkova, J. Thresholds of Replication Stress Signaling in Cancer Development and Treatment. Nat. Struct. Mol. Biol. 2012, 19, 5–7.

75. Tam, W.-L.; Ang, Y.-S.; Lim, B. The Molecular Basis of Ageing in Stem Cells. Mech. Ageing Dev. 2007, 128, 137–148.

76. Olovnikov, A.M. Telomeres, Telomerase, and Aging: Origin of the Theory. Exp. Gerontol. 1996, 31, 443–448.

77. Hayflick, L. THE LIMITED IN VITRO LIFETIME OF HUMAN DIPLOID CELL STRAINS. Exp. Cell Res. 1965, 37, 614–636.

78. Palacios, J.A.; Herranz, D.; De Bonis, M.L.; Velasco, S.; Serrano, M.; Blasco, M.A. SIRT1 Contributes to Telomere Maintenance and Augments Global Homologous Recombination. J. Cell Biol. 2010, 191, 1299–1313.

79. Rijo-Ferreira, F.; Takahashi, J.S. Genomics of Circadian Rhythms in Health and Disease. Genome Med. 2019, 11, 82.

80. Feillet, C.; van der Horst, G.T.J.; Levi, F.; Rand, D.A.; Delaunay, F. Coupling between the Circadian Clock and Cell Cycle Oscillators: Implication for Healthy Cells and Malignant Growth. Front. Neurol. 2015, 6, 96.

81. Langmesser, S.; Tallone, T.; Bordon, A.; Rusconi, S.; Albrecht, U. Interaction of Circadian Clock Proteins PER2 and CRY with BMAL1 and CLOCK. BMC Mol. Biol. 2008, 9, 41.

82. Takahashi, J.S. Transcriptional Architecture of the Mammalian Circadian Clock. Nat. Rev. Genet. 2017, 18, 164–179.

83. Yang, X. A Wheel of Time: The Circadian Clock, Nuclear Receptors, and Physiology. Genes Dev. 2010, 24, 741–747.

84. Stavreva, D.A.; Garcia, D.A.; Fettweis, G.; Gudla, P.R.; Zaki, G.F.; Soni, V.; McGowan, A.; Williams, G.; Huynh, A.; Palangat, M.; et al. Transcriptional Bursting and Co-Bursting Regulation by Steroid Hormone Release Pattern and Transcription Factor Mobility. Mol. Cell 2019, 75, 1161–1177.e11.

85. Yan, J.; Goldbeter, A. Robust Synchronization of the Cell Cycle and the Circadian Clock through Bidirectional Coupling. J. R. Soc. Interface 2019, 16, 20190376.

86. Kowalska, E.; Ripperger, J.A.; Hoegger, D.C.; Bruegger, P.; Buch, T.; Birchler, T.; Mueller, A.; Albrecht, U.; Contaldo, C.; Brown, S.A. NONO Couples the Circadian Clock to the Cell Cycle. Proc. Natl. Acad. Sci. U. S. A. 2013, 110, 1592–1599.

87. Umemura, Y.; Yagita, K. Development of the Circadian Core Machinery in Mammals. J. Mol. Biol. 2020, 432, 3611–3617.

88. Vallone, D.; Lahiri, K.; Dickmeis, T.; Foulkes, N.S. Start the Clock! Circadian Rhythms and Development. Dev. Dyn. 2007, 236, 142–155.

89. Raleigh, J.M.; O’Connell, M.J. The G(2) DNA Damage Checkpoint Targets Both Wee1 and Cdc25. J. Cell Sci. 2000, 113, 1727–1736.

90. Burchett, J.B.; Knudsen-Clark, A.M.; Altman, B.J. MYC Ran Up the Clock: The Complex Interplay between MYC and the Molecular Circadian Clock in Cancer. Int. J. Mol. Sci. 2021, 22, doi:10.3390/ijms22147761.

91. Gotoh, T.; Vila-Caballer, M.; Santos, C.S.; Liu, J.; Yang, J.; Finkielstein, C.V. The Circadian Factor Period 2 Modulates p53 Stability and Transcriptional Activity in Unstressed Cells. Mol. Biol. Cell 2014, 25, 3081–3093.

92. Stokes, K.; Nunes, M.; Trombley, C.; Flôres, D.E.F.L.; Wu, G.; Taleb, Z.; Alkhateeb, A.; Banskota, S.; Harris, C.; Love, O.P.; et al. The Circadian Clock Gene, Bmal1, Regulates Intestinal Stem Cell Signaling and Represses Tumor Initiation. Cell Mol Gastroenterol Hepatol 2021, 12, 1847–1872.e0.

93. Jiang, L.; Zhang, F.; Fan, W.; Zheng, M.; Kang, J.; Huang, F.; He, H. Expression of Circadian Clock Genes during Differentiation of Rat Dental Papilla Cells in Vitro. Biological Rhythm Research 2020, 1–12.

94. Morse, D.; Cermakian, N.; Brancorsini, S.; Parvinen, M.; Sassone-Corsi, P. No Circadian Rhythms in Testis: Period1 Expression Is Clock Independent and Developmentally Regulated in the Mouse. Mol. Endocrinol. 2003, 17, 141–151.

95. Beaver, L.M.; Rush, B.L.; Gvakharia, B.O.; Giebultowicz, J.M. Noncircadian Regulation and Function of Clock Genes Period and Timeless in Oogenesis of Drosophila Melanogaster. J. Biol. Rhythms 2003, 18, 463–472.

96. Bittman, E.L. Timing in the Testis. J. Biol. Rhythms 2016, 31, 12–36.

97. Lu, Y.; Zheng, X.; Hu, W.; Bian, S.; Zhang, Z.; Tao, D.; Liu, Y.; Ma, Y. Cancer/testis Antigen PIWIL2 Suppresses Circadian Rhythms by Regulating the Stability and Activity of BMAL1 and CLOCK. Oncotarget 2017, 8, 54913–54924.

98. Solov’eva, L.; Svetlova, M.; Bodinski, D.; Zalensky, A.O. Nature of Telomere Dimers and Chromosome Looping in Human Spermatozoa. Chromosome Res. 2004, 12, 817–823.

99. Walen, K.H. Neoplastic-like CELL Changes of Normal Fibroblast Cells Associated with Evolutionary Conserved Maternal and Paternal Genomic Autonomous Behavior (gonomery). J. Cancer Ther. 2014, 05, 860–877.

100. Mayer, W.; Smith, A.; Fundele, R.; Haaf, T. Spatial Separation of Parental Genomes in Preimplantation Mouse Embryos. J. Cell Biol. 2000, 148, 629–634.

101. Curran, K.L.; Allen, L.; Porter, B.B.; Dodge, J.; Lope, C.; Willadsen, G.; Fisher, R.; Johnson, N.; Campbell, E.; VonBergen, B.; et al. Circadian Genes, xBmal1 and xNocturnin, Modulate the Timing and Differentiation of Somites in Xenopus Laevis. PLoS One 2014, 9, e108266.

102. Bellet, M.M.; Orozco-Solis, R.; Sahar, S.; Eckel-Mahan, K.; Sassone-Corsi, P. The Time of Metabolism: NAD+, SIRT1, and the Circadian Clock. Cold Spring Harb. Symp. Quant. Biol. 2011, 76, 31–38.

103. Wang, R.-H.; Zhao, T.; Cui, K.; Hu, G.; Chen, Q.; Chen, W.; Wang, X.-W.; Soto-Gutierrez, A.; Zhao, K.; Deng, C.-X. Negative Reciprocal Regulation between Sirt1 and Per2 Modulates the Circadian Clock and Aging. Sci. Rep. 2016, 6, 28633.

104. Yang, X.; Wood, P.A.; Hrushesky, W.J.M. Mammalian TIMELESS Is Required for ATM-Dependent CHK2 Activation and G2/M Checkpoint Control. Journal of Biological Chemistry 2010, 285, 3030–3034.

105. Chao, H.-W.; Doi, M.; Fustin, J.-M.; Chen, H.; Murase, K.; Maeda, Y.; Hayashi, H.; Tanaka, R.; Sugawa, M.; Mizukuchi, N.; et al. Circadian Clock Regulates Hepatic Polyploidy by Modulating Mkp1-Erk1/2 Signaling Pathway. Nat. Commun. 2017, 8, 2238.

106. Bernstein, B.E.; Mikkelsen, T.S.; Xie, X.; Kamal, M.; Huebert, D.J.; Cuff, J.; Fry, B.; Meissner, A.; Wernig, M.; Plath, K.; et al. A Bivalent Chromatin Structure Marks Key Developmental Genes in Embryonic Stem Cells. Cell 2006, 125, 315–326.

107. Fung-Uceda, J.; Lee, K.; Seo, P.J.; Polyn, S.; De Veylder, L.; Mas, P. The Circadian Clock Sets the Time of DNA Replication Licensing to Regulate Growth in Arabidopsis. Dev. Cell 2018, 45, 101–113.e4.

108. Anatskaya, O.V.; Vinogradov, A.E.; Vainshelbaum, N.M.; Giuliani, A.; Erenpreisa, J. Phylostratic Shift of Whole-Genome Duplications in Normal Mammalian Tissues towards Unicellularity Is Driven by Developmental Bivalent Genes and Reveals a Link to Cancer. Int. J. Mol. Sci. 2020, 21, doi:10.3390/ijms21228759.

109. Vazquez-Martin, A.; Anatskaya, O.V.; Giuliani, A.; Erenpreisa, J.; Huang, S.; Salmina, K.; Inashkina, I.; Huna, A.; Nikolsky, N.N.; Vinogradov, A.E. Somatic Polyploidy Is Associated with the Upregulation of c-MYC Interacting Genes and EMT-like Signature. Oncotarget 2016, 7, 75235–75260.

110. Dakup, P.P.; Porter, K.I.; Gajula, R.P.; Goel, P.N.; Cheng, Z.; Gaddameedhi, S. The Circadian Clock Protects against Ionizing Radiation-Induced Cardiotoxicity. FASEB J. 2020, 34, 3347–3358.

111. Dakup, P.P.; Porter, K.I.; Gaddameedhi, S. The Circadian Clock Protects against Acute Radiation-Induced Dermatitis. Toxicol. Appl. Pharmacol. 2020, 399, 115040.

112. Shilts, J.; Chen, G.; Hughey, J.J. Evidence for Widespread Dysregulation of Circadian Clock Progression in Human Cancer. PeerJ 2018, 6, e4327.

113. Wu, Y.; Tao, B.; Zhang, T.; Fan, Y.; Mao, R. Pan-Cancer Analysis Reveals Disrupted Circadian Clock Associates With T Cell Exhaustion. Front. Immunol. 2019, 10, 2451.

114. Relógio, A.; Thomas, P.; Medina-Pérez, P.; Reischl, S.; Bervoets, S.; Gloc, E.; Riemer, P.; Mang-Fatehi, S.; Maier, B.; Schäfer, R.; et al. Ras-Mediated Deregulation of the Circadian Clock in Cancer. PLoS Genet. 2014, 10, e1004338.

115. Van Dycke, K.C.G.; Rodenburg, W.; van Oostrom, C.T.M.; van Kerkhof, L.W.M.; Pennings, J.L.A.; Roenneberg, T.; van Steeg, H.; van der Horst, G.T.J. Chronically Alternating Light Cycles Increase Breast Cancer Risk in Mice. Curr. Biol. 2015, 25, 1932–1937.

116. Papagiannakopoulos, T.; Bauer, M.R.; Davidson, S.M.; Heimann, M.; Subbaraj, L.; Bhutkar, A.; Bartlebaugh, J.; Vander Heiden, M.G.; Jacks, T. Circadian Rhythm Disruption Promotes Lung Tumorigenesis. Cell Metab. 2016, 24, 324–331.

117. Zhang, J.; Lv, H.; Ji, M.; Wang, Z.; Wu, W. Low Circadian Clock Genes Expression in Cancers: A Meta-Analysis of Its Association with Clinicopathological Features and Prognosis. PLoS One 2020, 15, e0233508.

118. Tomczak, K.; Czerwińska, P.; Wiznerowicz, M. The Cancer Genome Atlas (TCGA): An Immeasurable Source of Knowledge. Contemp. Oncol. 2015, 19, A68–A77.

119. Rahman, M.; Jackson, L.K.; Johnson, W.E.; Li, D.Y.; Bild, A.H.; Piccolo, S.R. Alternative Preprocessing of RNA-Sequencing Data in The Cancer Genome Atlas Leads to Improved Analysis Results. Bioinformatics 2015, 31, 3666–3672.

120. Lu, C.; Yang, Y.; Zhao, R.; Hua, B.; Xu, C.; Yan, Z.; Sun, N.; Qian, R. Role of Circadian Gene Clock during Differentiation of Mouse Pluripotent Stem Cells. Protein Cell 2016, 7, 820–832.

121. Carter, S.L.; Cibulskis, K.; Helman, E.; McKenna, A.; Shen, H.; Zack, T.; Laird, P.W.; Onofrio, R.C.; Winckler, W.; Weir, B.A.; et al. Absolute Quantification of Somatic DNA Alterations in Human Cancer. Nat. Biotechnol. 2012, 30, 413–421.

122. Taylor, A.M.; Shih, J.; Ha, G.; Gao, G.F.; Zhang, X.; Berger, A.C.; Schumacher, S.E.; Wang, C.; Hu, H.; Liu, J.; et al. Genomic and Functional Approaches to Understanding Cancer Aneuploidy. Cancer Cell 2018, 33, 676–689.e3.

123. Salmina; Salmina; Gerashchenko; Hausmann; Vainshelbaum; Zayakin; Erenpreiss; Freivalds; Cragg; Erenpreisa When Three Isn’t a Crowd: A Digyny Concept for Treatment-Resistant, Near-Triploid Human Cancers. Genes 2019, 10, 551.

124. Vainshelbaum, N.M.; Zayakin, P.; Kleina, R.; Giuliani, A.; Erenpreisa, J. Meta-Analysis of Cancer Triploidy: Rearrangements of Genome Complements in Male Human Tumors Are Characterized by XXY Karyotypes. Genes 2019, 10, doi:10.3390/genes10080613.

125. Vinnitsky, V.B. Oncogerminative Hypothesis of Tumor Formation. Med. Hypotheses 1993, 40, 19–27.

126. Grundmann, E. [The concept of Julius Cohnheim on tumor formation and metastasis from the viewpoint of new research results]. Zentralbl. Allg. Pathol. 1985, 130, 323–331.

127. Lange, U.C.; Saitou, M.; Western, P.S.; Barton, S.C.; Surani, M.A. The Fragilis Interferon-Inducible Gene Family of Transmembrane Proteins Is Associated with Germ Cell Specification in Mice. BMC Dev. Biol. 2003, 3, 1.

128. Borghesan, M.; Fafián-Labora, J.; Eleftheriadou, O.; Carpintero-Fernández, P.; Paez-Ribes, M.; Vizcay-Barrena, G.; Swisa, A.; Kolodkin-Gal, D.; Ximénez-Embún, P.; Lowe, R.; et al. Small Extracellular Vesicles Are Key Regulators of Non-Cell Autonomous Intercellular Communication in Senescence via the Interferon Protein IFITM3. Cell Rep. 2019, 27, 3956–3971.e6.

129. Sheng, X.; Tian, C.; Liu, L.; Wang, L.; Ye, X.; Li, J.; Zeng, M.; Liu, L. Characterization of Oogonia Stem Cells in Mice by Fragilis. Protein Cell 2019, 10, 825–831.

130. Bruggeman, J.W.; Irie, N.; Lodder, P.; van Pelt, A.M.M.; Koster, J.; Hamer, G. Tumors Widely Express Hundreds of Embryonic Germline Genes. Cancers 2020, 12, doi:10.3390/cancers12123812.

131. Pjanova, D.; Vainshelbaum, N.M.; Salmina, K.; Erenpreisa, J. The Role of the Meiotic Component in Reproduction of B-RAF-Mutated Melanoma: A Review and “Brainstorming” Session. Melanoma 2021.

